# Cell-free chromatin particles released from dying cells inflict mitochondrial damage and ROS production in living cells

**DOI:** 10.1101/2021.12.30.474529

**Authors:** Gorantla V Raghuram, Bhabesh Kumar Tripathy, Kartikeya Avadhani, Snehal Shabrish, Naveen Kumar Khare, Relestina Lopes, Kavita Pal, Indraneel Mittra

## Abstract

mtDNA damage and the resultant oxidative stress are associated with neurodegenerative diseases, ageing and cancer. However, what triggers mtDNA damage remains unclear. We have reported that cell-free chromatin particles (cfChPs) that are released from the billions of cells that die in the body every day can readily enter into healthy cells and damage their DNA. We show here that cfChPs isolated from sera of healthy individuals, or those that are released from dying cells, inflict direct physical damage mtDNA leading to marked activation of ROS. The latter could be abrogated by concurrent treatment with three different cfChPs deactivating agents. Given that 1×10^9^-1×10^12^ cells die in the body every day, our findings suggest that cfChPs from dying cells are major physiological triggers for mtDNA damage and ROS production. Deactivation of cfChPs may provide a novel therapeutic approach to retard ageing and associated degenerative conditions that have been linked to oxidative stress.

## Introduction

Mitochondrial electron transport chain during oxidative phosphorylation is thought to be the predominant source of intracellular ROS^1^. In addition, external factors such as UV radiation, chemicals, heat and pH stress can also lead to production of mitochondrial ROS^2–5^. Under normal physiological conditions, ROS levels are regulated by a balance between generation of ROS and their elimination by several antioxidant enzymes and non-enzymatic defence systems^6^. Oxidative stress can arise when there is excess ROS production or its inefficient elimination. One of the major triggers of excess ROS production is damage to mitochondria, especially to mtDNA^7, 8^. Excess ROS production can cause irreversible damage to mtDNA, proteins and membrane lipids resulting in mitochondrial dysfunction and ultimately to cell death^9^. Chronic mitochondrial dysfunction is associated with ageing and several degenerative conditions such as diabetes, cardiovascular diseases, neurodegenerative disorders and cancer^10–14^.

Several hundred billion to a trillion cells die in the body every day^15,16^ and the cell-free chromatin particles (cfChPs) that are released from them enter into the extracellular compartments, including into the circulation^17^. We have earlier reported that cfChPs can readily enter into the cells and damage their DNA^18, 19^. In this study we provide evidence that cfChPs are a major source of ROS production by their ability to readily enter into healthy cells and inflict damage to mitochondria, especially to mtDNA. We also provide evidence that mtDNA damage and ROS production can be minimized by treatment with cfChPs inactivating agents suggesting therapeutic possibilities for multiple conditions mentioned above that are associated with oxidative stress.

## Results

### cfChPs are readily internalized by NIH3T3 cells

cfChPs isolated from sera of healthy individuals were fluorescently dually labelled in their DNA with Platinum Bright 550 and in their histones with ATTO-488 and added to cultured NIH3T3 cells (10 ng equivalent of DNA). Abundant fluorescent signals were detected in the cytoplasm of the treated cells at 4 h (**Fig. 1**). All further experiments described below were conducted at 4 h.

**Fig. 1:**
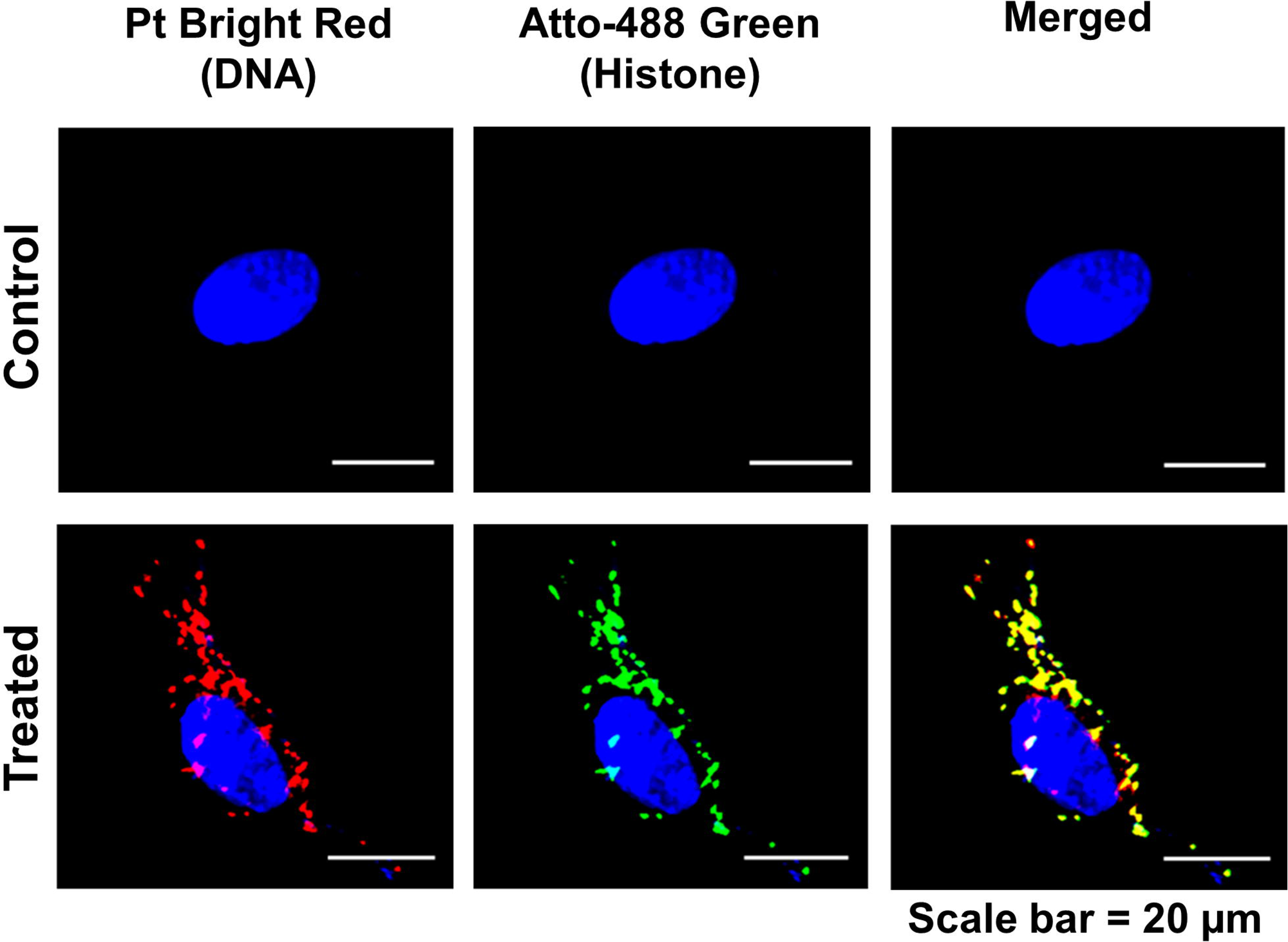
Rapid uptake of cfChPs by NIH3T3 cells at 4h. NIH3T3 cells were treated with cfChPs (10ng) isolated from sera of healthy individuals which had been dually labelled in their DNA (Platinum Bright 550) and in their histones (ATTO-488). Images were acquired on Applied Spectral bio-imaging system. Dually labelled cfChPs are clearly seen in the cytoplasm. The particles appear relatively large which is most likely due to linking-up of multiple cfChPs to form concatameres as described by us earlier^18^.

### cfChPs inflict mtDNA damage: studies on whole cells

In order to investigate whether the internalized cfChPs induced mtDNA damage, NIH3T3 cells were treated with cfChPs (10ng) and the treated cells were immune stained with antibodies against γH2AX and pATM. The cells were simultaneously stained with MitoTracker Red to identify mitochondria. Figure 2 shows that numerous fluorescence γH2AX and pATM signals could be detected in the cytoplasm of treated cells which were highly significantly higher than those detected in untreated control cells (p<0.0001) **(Fig. 2)**.

**Fig. 2.**
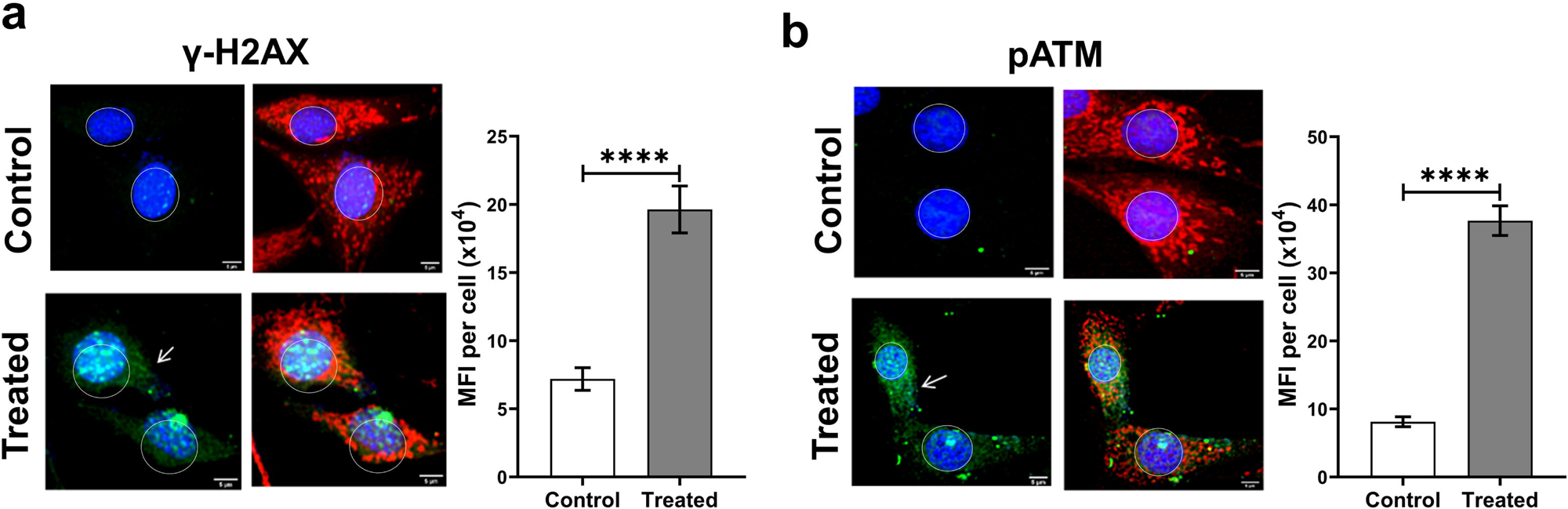
cfChPs treatment of NIH3T3 cells induces mitochondrial DNA damage as observed at 4h. **a.** Representative images of untreated control and cfChPs (10ng) treated cells dually stained with MitoTracker red dye and antibody against γH2AX (green). Expression of γH2AX is visibly increased in the mitochondria of cfChPs treated cells (arrows). Quantitative analysis of MFI was performed after gating and excluding the nuclear fluorescence. Analysis illustrates significantly higher MFI for γH2AX in treated cells as compared to untreated controls. It should be noted that cfChPs have induced the expected activation of H2AX in the nuclei of the treated cells as reported by us earlier^18^**. b.** Representative images of untreated control and cfChPs (10ng) treated cells dually stained with MitoTracker red dye and antibody against pATM (green). Expression of pATM is visibly increased in the mitochondria of cfChPs treated cells (arrows). Quantitative analysis of MFI was performed after gating and excluding the nuclear fluorescence. Analysis illustrates significantly higher MFI for pATM in treated cells as compared to untreated controls. It should be noted that cfChPs have induced the expected activation of ATM in the nuclei of the treated cells as reported by us earlier^18^. Experiments were repeated twice and all images were acquired under the same settings. Results represent mean ± SEM values. Scale Bar - 5 µm. Data were analysed using Student’s_t-_test. *** p<0.001, **** p<0.0001.

### cfChPs inflict mtDNA damage: studies using isolated mitochondria

Mitochondria were isolated from NIH3T3 cells that had been treated with cfChPs (10ng) for 4h. Isolated mitochondria were clearly identifiable as DAPI positive particles which co-localized with MitoTracker Red **(Fig. 3a)**. The isolated mitochondria were next analysed in immune FISH experiments using fluorescent antibodies against histone H4 and a human whole genomic DNA probe. Negative control experiments, not shown here, were done to confirm that the DNA FISH probe did not cross hybridized with mouse DNA. Strictly co-localising fluorescent signals of histone H4, human DNA with DAPI indicated that cfChPs were in close contact with mouse mitochondria represented by DAPI positive signals **(Fig. 3b)**. Dual staining of the isolated mitochondria with antibodies against γH2AX and MitoTracker Red and pATM and MitoTracker Red detected strictly co-localising signals indicating that cfChPs that were associated with mitochondria had inflicted mtDNA damage **(Figs 3c and d)**. When compared with isolated mitochondria from untreated NIH3T3 cells, the number of γH2AX and pATM signals in mitochondria isolated from treated cells were found to be significantly elevated (p<0.001 and p<0.0001 respectively).

**Fig. 3.**
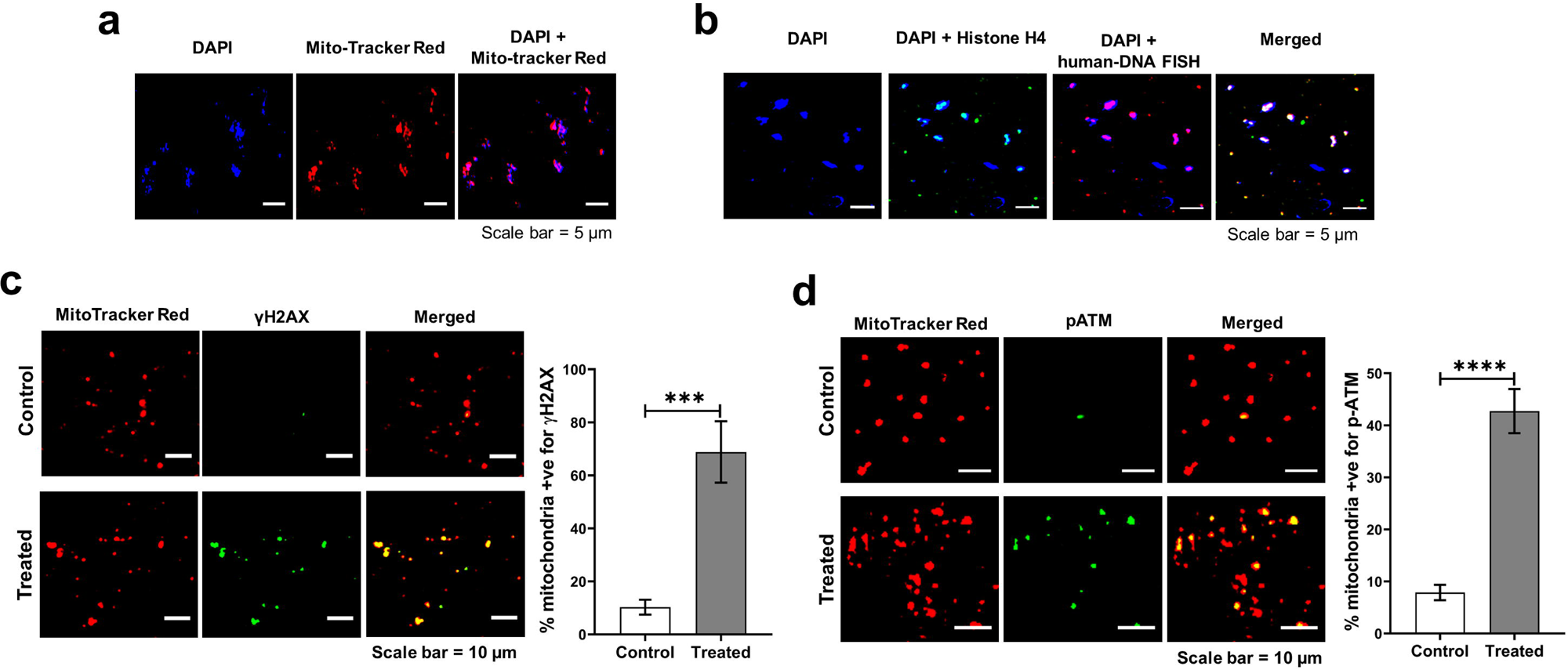
Mitochondria isolated from cfChPs treated cells show intimate association with cfChPs. **a.** Confirmation that isolated mitochondria represented by Mito-Tracker Red co-localize with DAPI signals; **b.** Immuno-FISH performed on isolated mitochondria using antibody against histone H4 and a human DNA FISH probe show co-localization of cfChPs with mitochondria represented by DAPI. **c.** Mitochondria isolated from cfChPs treated cells show phosphorylated H2AX signals which co-localize with mitochondria represented by Mito-Tracker Red. Few γH2AX signals are seen in untreated control cells. Histograms represent quantitative analysis of co-localizing signals in control and treated cells. **d.** Mitochondria isolated from cfChPs treated cells show phosphorylated ATM signals which co-localize with mitochondria represented by Mito-Tracker Red. Few pATM signals are seen in untreated control cells. Histograms represent quantitative analysis of co-localizing signals in control and treated cells. Results are represented as mean ± SEM values and data were analysed using Student’s_t*-*_test. **\*\*\*** p<0.001; **\*\*\*\*** p<0.0001.

### cfChPs damage other components of mitochondria

We next examined whether, in addition to mtDNA damage, cfChPs inducing damage to other components of mitochondria.

#### Ultra structural changes

When examined under transmission electron microscopy (TEM), the untreated control cells were found to harbour mitochondria that were normal and elongated in shape. Mitochondria of the cfChPs treated cells, in contrast, were mainly rounded in appearance. The latter has been reported to signify mitochondrial damage^20^ (**Fig. 4a**). Proportion of elongated and round mitochondria in control and treated cells are depicted as histograms. Quantitative estimation of the ratio of round versus elongated mitochondria showed a three-fold increase in favour of round mitochondria indicating increased mitochondrial damage in the treated cells (**Fig. 4a**).

**Fig. 4:**
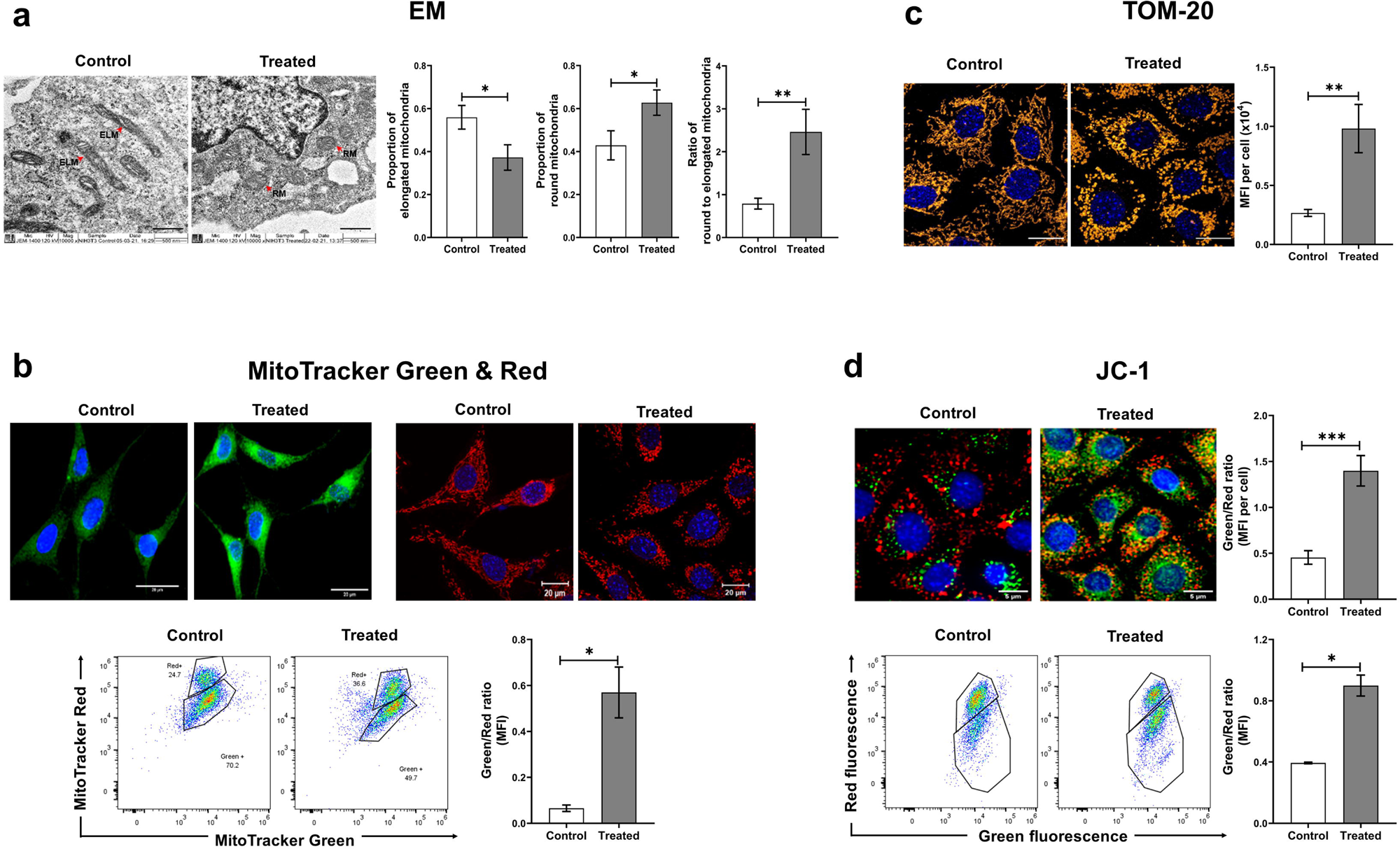
Various mitochondrial abnormalities induced by cfChPs in treated cells as observed at 4 h. **a.** Representative EM images of untreated and cfChPs (10ng) treated cells reveals ultra-structural changes in mitochondria. Control cells contain elongated mitochondria (red arrows), whereas cfChPs treated cells harbour mostly round shaped mitochondria (red arrows) (Scale bar - 1 µm). ELM= Elongated Mitochondria, RM= Round Mitochondria. Histograms represent quantitative analysis of ultrastructural changes and the ratio of round to elongated mitochondria in treated and control cells. **b.** Evaluation of mitochondrial dysfunction by IF and flow cytometry using mitochondrial dyes. Representative microphotographs of mitochondrial mass (MitoTracker Green) and of mitochondrial dysfunction (MitoTracker red) in cfChPs (10ng) treated and control cells. Quantitative analysis (MFI) of the ratio of mitochondrial mass and mitochondrial dysfunction by flow cytometry. The latter shows a significant increase in ratio of green to red fluorescence in cfChPs treated cells as compared to untreated cells. **c.** Representative confocal microphotographs of mitochondrial outer membrane marker TOM20 in cfChPs treated (10ng) and untreated cells. Expression of TOM20 is visibly higher in cfChPs treated cells compared to untreated cells. (Scale Bar - 20 µm, Pseudo colour - Orange). Quantitative analysis (MFI) of TOM20 expression in control versus cfChPs treated cells exhibits a significant increase in MFI in treated cells indicating loss of mitochondrial membrane integrity as compared to control cells. **d.** Representative confocal microphotographs of mitochondrial membrane potential marker JC-1 in cfChPs treated (10ng) and untreated cells. Quantitative analysis (MFI) shows increased green to red fluorescence ratio signifying depolarized mitochondrial membrane. Quantitative MFI analysis by flow cytometry reveals higher ratio of green to red fluorescence in cfChPs treated cells compared to control cells. All the experiments were repeated twice. Results are represented as mean ± SEM values and data were analysed using Student’s_t*-*_test. * =p<0.05, ** =p<0.01, *** =p<0.005.

#### Increase in mitochondrial mass and mitochondrial shape

It has been reported that a cell up-regulates mitochondrial biogenesis in response to mitochondrial damage in the form of increased mitochondrial mass^21^. MitoTracker Green has been widely used to assess mitochondrial mass as it stains the mitochondria irrespective of its membrane potential^22^. A marked increase in MitoTracker Green fluorescence was observed in cells treated with cfChPs indicating that mitochondrial mass had increased as a result of damage inflicted by cfChPs treatment (**Fig. 4b**). MitoTracker red is known to detect changes in mitochondrial shape which correlates with mitochondrial damage^20, 23^. Confocal microscopy of cfChPs treated cells stained with MitoTracker Red generated diffuse fluorescence signals which have been reported to represent mitochondrial fragmentation^20^. Untreated control cells on the other hand exhibited a distinct network of mitochondria (**Fig. 4b).** Flow cytometric analysis of cells dually labelled with MitoTracker Green and Red was performed to determine the relative propotion of functional mitochondria (MitoTracker Green**^high^** and MitoTracker Red**^high^**) and dysfunctional mitochondria (MitoTracker Green**^high^** and MitoTracker Red**^low^**) in response to cfChPs treatment. Quantitative analysis revealed an increased ratio of MitoTracker Green**^high^** and MitoTracker Red**^low^** population in cfChPs treated cells compared to untreated cells. These findings provided additional evidence of mitochondrial dysfunction (**Fig. 4b**)

#### Upregulation of mitochondrial outer membrane protein TOM20

Translocase of the outer membrane 20 (TOM20) acts as an import receptor that belongs to the family of TOM proteins and allows movement of proteins across the mitochondrial outer membrane into the inner membrane space^24^. Upregulation of TOM20 is a sign of dysfunctional mitochondria with increased biogenesis and excess import of proteins^25^. We observed a significant increase in expression of TOM20 in cfChPs treated cells when compared to controls under confocal microscopy (**Fig. 4c**). Quantitative analysis expressed as MFI per cell showed a three-fold increase in TOM20 expression in cfChPs treated cells when compared to control untreated cells (p<0.01) (**Fig. 4c**).

#### Altered mitochondrial membrane potential

Low mitochondrial membrane potential has been reported to be associated with mitochondrial damage^26^. We investigated mitochondrial depolarization as an indicator of mitochondrial damage using JC-1 (5, 5’, 6, 6’-tetrachloro-1, 1’, 3, 3’-tetraethyl benzimidazolylcarbocyanine iodide) - a mitochondrial membrane fluorescent dye. JC-1 accumulates as aggregates in healthy mitochondria which are known to have a high membrane potential and emit red fluorescence. On the other hand, in dysfunctional mitochondria, JC-1 appears as monomers and emits green fluorescence since they have a relatively low membrane potential^27^. Representative images given in **Fig. 4d** show that while control cells have predominantly red signals, cfChPs treated cells show abundance of green signals in addition to red signals. Quantitative analysis of the data revealed a highly significant increase in ratio of green to red fluorescence signals in cells treated with cfChPs (p<0.001) (**Fig. 4d)**. These results were validated using flow cytometry which revealed a two-fold increase in ratio of green to red fluorescence in cfChPs treated cells compared to untreated cells (p<0.05). Taken together, these results indicate that cfChPs had induced damage to the mitochondria of the treated cells, leading to decreased mitochondrial membrane potential.

### cfChPs induce ROS production from mitochondria

Mitochondrial damage, especially mtDNA damage, is a major cause of excess ROS production^7, 8^. One of the primary forms of ROS generated in the mitochondria are superoxide radicals^9^, which can be readily detected by MitoSOX Red, a triphenylphosphonium (TPP+)-linked DHE compound that is selectively oxidized by superoxide radicals leading to emission of red fluorescence^28^. To investigate whether mitochondrial damage, including mtDNA damage that we have described above had led to increased ROS production, NIH3T3 cells were treated with cfChPs (10ng) for 4h followed by staining of the cells with MitoSOX Red. A marked increase in MitoSOX fluorescence was detected upon microscopic examination (p<0.0001) (**Fig. 5a)**. These results were validated using flow cytometry which detected a two-fold increase in MFI of MitoSox Red in cfChPs treated cells compared to untreated cells (p< 0.05). These data clearly indicated that cfChPs can induce increased ROS production resulting from damage to mitochondria, especially damage to mtDNA.

**Fig. 5:**
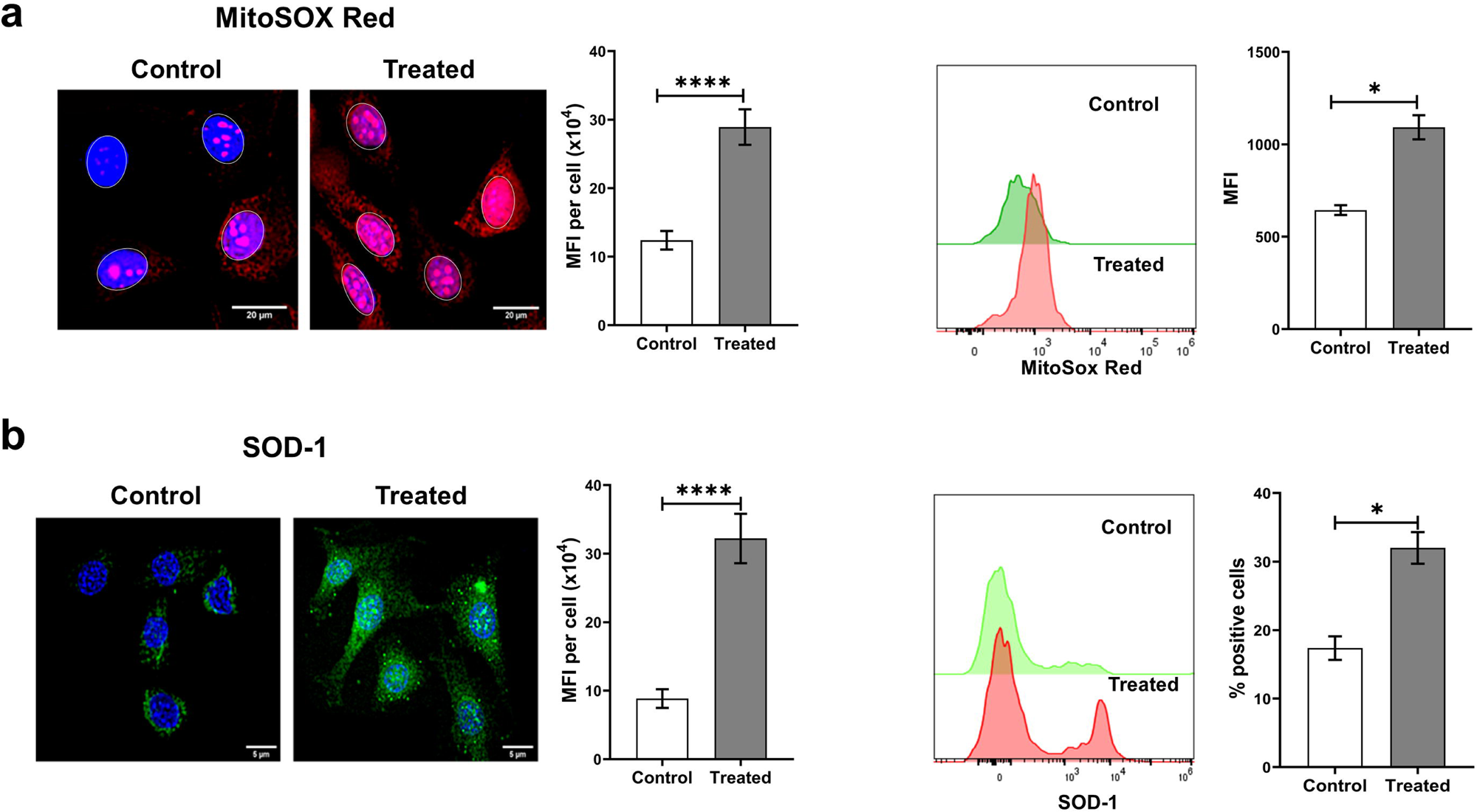
Treatment of NIH3T3 cells with cfChPs induces oxidative stress as detected at 4h. **a.** Representative fluorescence microscopy images of control and cfChPs treated (10ng) cells stained with MitoSOX Red shows significant activation of ROS production in treated cells as compared to untreated cells (Scale Bar = 5 µm). Results of quantitative analysis of MFI (after gating the nuclei to exclude nuclear fluorescence) are shown as histograms. Flow cytometric analysis of MitoSox Red confirms significant increase in treated cells compared to control cells. **b.** Representative fluorescence microscopy images of control and cfChPs (10ng) treated cells stained with superoxide dismutase-1 (SOD-1) antibody shows significant increase in activation of SOD-1 in treated cells as compared to untreated cells (Scale Bar = 5 µm). Results of quantitative analysis (MFI) are shown as histograms. Flow cytometric analysis of SOD-1 confirms significant increase in treated cells compared to control cells. All the experiments were repeated twice. Results are represented as mean ± SEM values and data were analysed using Student’s_t*-*_test. * =p<0.05, *** =p<0.005.

### cfChPs activate the antioxidant enzyme superoxide dismutase

The next question that we addressed was whether excess ROS production following treatment of NIH3T3 cells with cfChPs had led to an increase in the antioxidant enzyme SOD-1. The latter was investigated both by IF and flow cytometry. Fluorescence microscopy of cells treated with cfChPs (10ng) for 4h detected a highly significant increase in SOD-1 production (p<0.0001) (**Fig. 5b)**. Flow cytometry analysis which also showed increased SOD-1 expression in treated cells (p<0.05).

### Experiments using conditioned media from dying NIH3T3 cells

We next conducted experiments based on our earlier report that cfChPs that are spontaneously released from dying cells can be readily internalised by healthy cells^19^. These experiments involved treating NIH3T3 cells with conditioned medium containing cfChPs released from hypoxia induced dying NIH3T3 cells. The method for generating cfChPs containing conditioned medium has been described in the Material and Methods section. Supplementary figure 1 shows that cfChPs that were released from dually fluorescently labelled dying NIH3T3 cells were readily internalised by recipient healthy NIH3T3 cells. Conditioned medium containing cfChPs released from hypoxia induced dying NIH3T3 cells were collected and applied to live NIH3T3 cells. As a first step, we performed a dose response experiment wherein we added increasing volumes of cfChPs containing conditioned medium to NIH3T3 cells followed by assay for MitoSOX Red expression as a marker of ROS production. Since MitoSox Red generates the by-product mitoethidium which intercalates in DNA and generates nuclear fluorescence, nuclei were gated and nuclear fluorescence was excluded from MFI analysis. Optimum ROS production was seen with a volume of 50 µl of culture media (**Supplementary Fig. 2**). All further experiments were performed using 50µl of conditioned medium. In order to confirm that cfChPs in the conditioned medium were indeed responsible for ROS production, conditioned media were pre-treated with the three different cfChPs-deactivating agent viz. anti-histone antibody complexed nanoparticles (CNPs), DNase I and a novel pro-oxidant combination of resveratrol and copper (R-Cu). As noted earlier, 50 µl of conditioned medium induced a marked activation of ROS in NIH3T3 cells which was abrogated by pre-treatment of the conditioned medium with all three cfChPs inactivating agents (p<0.0001) **(Fig. 6a)**. This experiment was repeated using flow cytometry which also detected down regulation of ROS production by cfChPs inhibitors **(Fig. 6b).** It should be pointed out that the degree of reduction detected by flow cytometry was somewhat less than that of obtained by analysis by fluorescent microscopy, since the nuclear fluorescence could not be gated and excluded from analysis in flow-cytometry experiments.

**Fig. 6.**
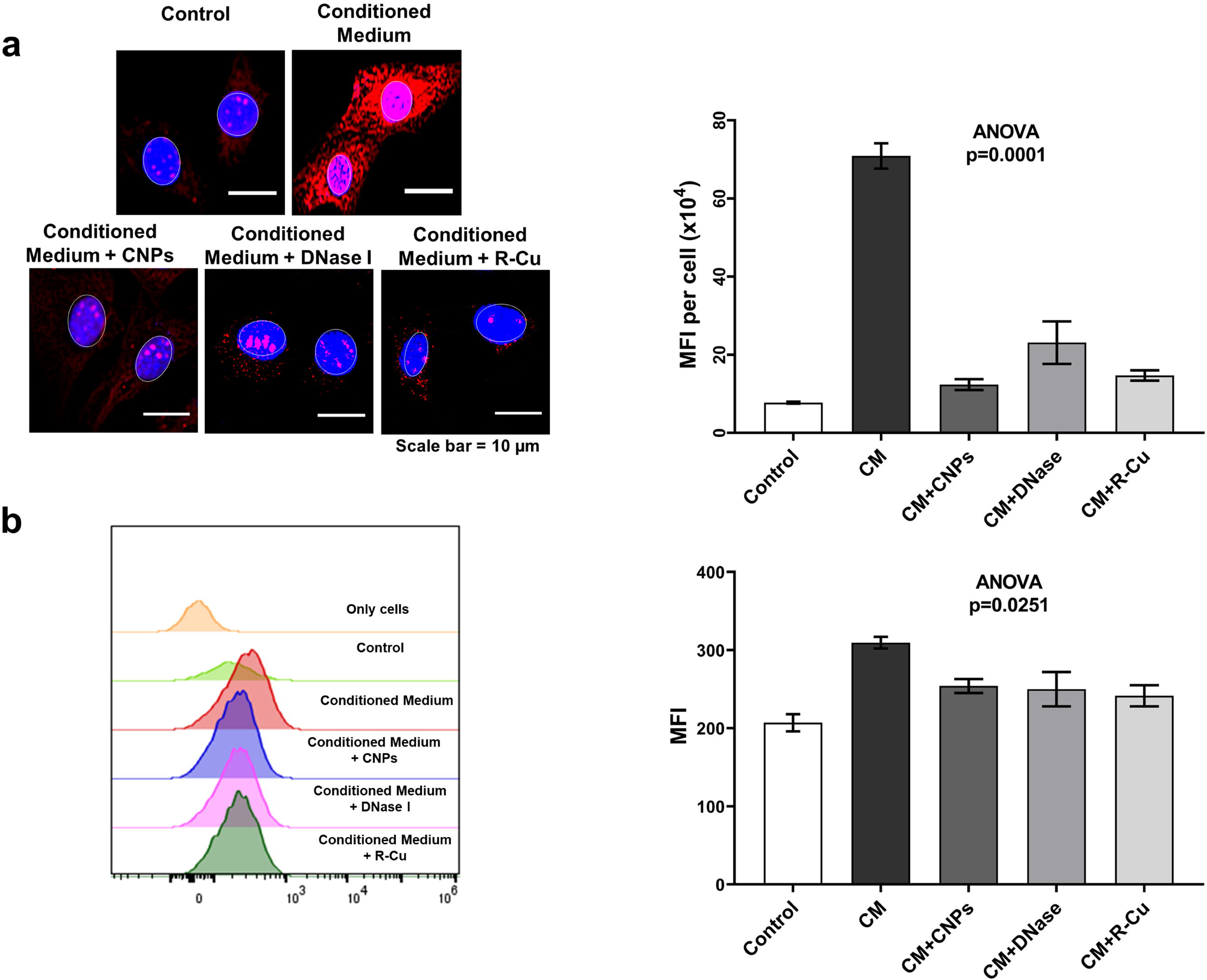
Activation of ROS production in NIH3T3 cells treated with conditioned medium containing cfChPs from hypoxia induced dying NIH3T3 cells in presence and absence of inhibitors of cfChPs. **a.** Representative fluorescence microscopy images showing activation of ROS production at 4h detected by MitoSox Red in NIH3T3 cells following treatment with conditioned medium (50µl) from dying NIH3T3 cells (upper panel). ROS activation is inhibited by concurrent treatment with CNPs, DNase I and R-Cu (lower panel). Results of quantitative analysis (MFI) are given as histograms. Since MitoSox Red generates the by-product mitoethidium which intercalates in DNA and generates nuclear fluorescence, nuclei were gated and nuclear fluorescence was excluded from MFI analysis. **b.** Representative flow cytometry plots showing activation of ROS production represented by MitoSox Red in NIH3T3 cells following treatment with conditioned medium (50µl) from dying NIH3T3 cells. ROS activation is inhibited by concurrent treatment with CNPs, DNase I and R-Cu. Results of quantitative analysis (MFI) are given as histograms. It should be noted that the degree of reduction detected by flow cytometry is less than that of obtained by microscopy analysis since the nuclear fluorescence could not be gated and excluded from analysis in flow-cytometry experiments. Experiments were repeated twice. Results are represented as mean ± SEM values and data were analysed using one way ANNOVA (GraphPad Prism 8).

## Discussion

Several hundred billion to a trillion cells die in the body every day^15, 16^, and cfChPs that are released from the dying cells enter into the extra-cellular compartments, including into the circulation^17^. We have earlier reported that circulating cfChPs, or those that are released locally from dying cells, can readily enter into healthy cells wherein they induce dsDNA breaks, inflammation and apoptotic responses^18, 19^. The current study was based on the hypothesis that, in addition to DNA damage, the internalised cfChPs would also induce damage to mitochondria. We have shown in this study that this is indeed the case. Treatment of NIH3T3 mouse fibroblast cells with cfChPs isolated from healthy human individuals led to activation of H2AX and ATM signals which co-localized with those of mitochondria identified by staining with MitoTracker Red. This was further confirmed in experiments using isolated mitochondria from cfChPs treated NIH3T3 cells in which we show that mtDNA damage is caused by physical association of cfChPs to mtDNA. We speculate that the physical association of cfChPs with mitochondria is the consequence of a chemical interaction between the positively charged histones of cfChPs and the negatively charged mtDNA. The binding of cfChPs to mitochondria leads to multiple other signs of mitochondrial damage including ultra-structural changes, changes in mitochondrial mass, shape and mitochondrial function. These events lead to activation of ROS production and consequent up-regulation of the anti-oxidant enzyme SOD-1. We propose that excess ROS production sets in motion a vicious cycle of more mtDNA damage and production of ROS leading to oxidative stress which is associated with several pathological conditions such as neurodegenerative diseases, ageing and cancer^7, 29, 30^ The mechanism of cfChPs induced mitochondrial damage and ROS production as discussed above is graphically illustrated in **Supplementary Fig. 3**.

We demonstrate that cfChPs induced ROS production can be greatly reduced by three different cfChPs inactivating agents. Of the three agents, a combination of the widely used nutraceuticals resveratrol (R) and copper (Cu) hold therapeutic promise. We have shown that combining small quantities of R and Cu leads to production of oxygen radicals which are capable of deactivating cfChPs with multiple therapeutic effects^31–38^.

Oxidative stress resulting from increased ROS production is known to be the underlying cause of many serious disorders^10–14^. However, what causes excess ROS production has remained elusive till now. We have shown in this study that mitochondrial damage is inflicted by extraneous agents in the form of cfChPs. Given that 1×10^9^-1×10^12^ cells die in the body every day, our findings suggest that cfChPs from dying cells are major physiological triggers for mtDNA damage and ROS production. Deactivation of cfChPs may provide a novel therapeutic approach to retard ageing and associated degenerative conditions that have been linked to oxidative stress. In this context, we have recently reported that prolonged administration of resveratrol and copper can retard multiple biological hallmarks of ageing and neurodegeneration in C57Bl/6 mice^38^.

## Materials and Methods

### Ethics approval

Institutional Ethics Committee (IEC) approval was obtained for collecting blood samples from healthy volunteers and signed informed consent was obtained on consent forms approved by IEC (Approval no.900520).

### Reagents and antibodies

Commercial sources and catalogue numbers of reagents and antibodies used in this study are given in Supplementary Table 1.

### Isolation of cfChPs from sera of healthy donors

Isolation of cfChPs from sera of healthy individuals was performed according to a protocol described by us earlier^18^. The isolated cfChPs when examined under electron microscopy revealed a beads-on-a-string appearance typical of chromatin^18^. The amount of DNA in cfChPs was estimated by PicoGreen quantification assay, and cfChPs concentration was expressed in terms of their DNA content.

### Fluorescent dual labelling of cfChPs

cfChPs were fluorescently dually labelled in their DNA by Platinum Bright 550 (red) and in their histone H4 with ATTO-488 (green) according to a protocol described by us earlier^18^.

### Cell culture

NIH3T3 embryonic mouse fibroblast cells were obtained from the American Type Culture Collection (ATCC, USA) and grown in Dulbecco’s Modified Eagle’s Medium (DMEM) (Gibco, Catalog No.12800-017) containing 10% bovine calf serum (Cytiva HyClone, Catalog No. SH30073) and maintained at 37_°C in an atmosphere of 5% CO_2_ and air. For microscopy experiments, cells were seeded at a density of 1×10^5^ on coverslips in 1.5 ml DMEM; for flow cytometry, 2×10^5^ cells were seeded in 1.5 ml DMEM and incubated overnight before conducting the experiments.

### Fluorescence and confocal microscopy

For fluorescence microscopy cells were imaged using Spectral Bio-Imaging System (Applied Spectral Imaging). Images were captured using a 40x air objective. All images were captured with the same exposure time set to ensure uniform fluorescence intensity for purposes of comparison. For confocal microscopy, images were acquired using a 63x oil objective on Zeiss LSM 780 laser scanning microscope (Carl Zeiss, Germany) and Leica SP8 confocal imaging system (Leica Microsystems, Wetzlar, Germany). Mean fluorescence intensity (MFI) of images was measured using ImageJ (W. Rasband, Bethesda, USA).

### Flow cytometry

For flow cytometry, cells were acquired on Attune NxT flow cytometer (Thermo Fisher Scientific, USA) and analysis was performed using *FlowJo*™ *v10.6 Software*. NIH3T3 cells were gated on forward and side scatter parameters. At least 20,000 NIH3T3 cells were acquired and the threshold between negative and positive expression was defined by the fluorescence minus one (FMO) method. Median fluorescence intensity (MFI) of respective dye was then compared by histogram analysis of untreated and treated cells.

### Isolation of mitochondria

NIH3T3 cells were treated with 10 ng cfChPs for 4 h and mitochondria were isolated from the treated cells using Mitochondria Isolation Kit (Supplementary Table 1) as per the instruction manual. Mitochondria were similarly isolated from control untreated cells.

### Fluorescent staining of isolated mitochondria

#### a) MitoTracker Red CMXRos Staining

Isolated mitochondria were cytospun onto a slide and fixed in 2% paraformaldehyde (PFA) / 1.5% glutaraldehyde (GA) fixative mixture for 15 min on ice followed by neutralisation in 0.3 M Glycine buffer for 30 min at room temperature. The fixed smears were stained with 100nM MitoTracker Red CMXRos for 15 min at room temperature and stained with DAPI (1µg/ml) in Hanks Balanced Salt Solution (HBSS). Images were acquired on the Spectral Bio-Imaging System (Applied Spectral Imaging).

#### b) Immuno-FISH for Histone and Human DNA

Isolated mitochondria were fixed as described above and processed for immune-staining with anti-histone H4 IgG and appropriate secondary antibodies (Supplementary Table 1). The immunostained smears were fixed with 2% PFA - 1.5% GA mixture for 5 min on ice and processed for FISH using custom synthesized Human DNA probe (Supplementary Table 1). Briefly, immune-stained slides were hybridised overnight with Human DNA probe at 37°C and washed once with 0.4X sodium saline Citrate (SSC) at 72°C (± 2°C) for 2 mins and 4X SSC in Tween-20 twice at room temperature for 5 mins each. The slides were mounted with DAPI (1µg/ml) in HBSS and images were acquired on the Spectral Bio-Imaging System (Applied Spectral Imaging).

### Detection of mtDNA damage by immunofluorescence

Mitochondrial DNA damage in the form of activation of H2AX and ATM was evaluated on both whole cells and isolated mitochondria.

#### Whole Cells

Cells were stained with the mitochondrial stain MitoTracker Red CMXRos and processed for immune-staining for γ-H2AX and p-ATM using appropriate antibodies (Supplementary table 1). The cells were counterstained with Vecta-shield DAPI and mounted on slides. Images were acquired using Spectral Bio-Imaging System (Applied Spectral Imaging). Mean fluorescence intensity (MFI) in the cytoplasm was measured after gating out the nuclei fluorescence.

#### Isolated mitochondria

DNA damage in isolated mitochondria were assessed by standard IF procedure as described above. Images were acquired under Spectral Bio-Imaging System (Applied Spectral Imaging).

### Assessment of mitochondrial integrity by electron microscopy

Exponentially growing NIH3T3 cells (4×10^6^) were treated with cfChPs (10 ng) for 4 h. Control and treated cells were fixed with 3% glutaraldehyde in 0.1 M sodium cacodylate buffer (pH 7.2) for 2 h at 4°C followed by 1% osmium tetroxide in 0.1 M sodium cacodylate buffer for 1 h at 4°C. Samples were then dehydrated using alcohol at 4°C and embedded in araldite resin to make ultrathin sections that were mounted onto EM grids for imaging. Grids were examined under a JEM1400-Plus transmission electron microscope (JEOL, Japan).

### Assessment of mitochondrial mass using MitoTracker Green FM

#### Fluorescence microscopy

Following treatment with cfChPs (10 ng) for 4 h, NIH3T3 cells were stained with 100 nM MitoTracker Green FM diluted in 2ml of Hank’s balanced salt solution (HBSS, pH 7.4) for 15 minutes at 37°C. Cells were washed with HBSS, counterstained with Hoechst, mounted on glass slides and imaged using Spectral Bio-Imaging System.

#### Flow cytometry

Cells were stained with 20nM MitoTracker Green FM diluted in 2ml of Hank’s balanced salt solution (HBSS, pH 7.4) for 15 minutes at 37°C. Cells were washed twice with pre-warm HBSS, collected in FACs tubes using cell scraper. Samples were acquired on Attune NxT flow cytometer (ThermoFisher Scientific, USA)

### Assessment of mitochondrial shape using MitoTracker Red CMXRos

#### Fluorescence microscopy

Following treatment with cfChPs (10 ng) for 4 h, NIH3T3 cells were stained with MitoTracker Red CMXRos (100nM) diluted in 2ml of Hank’s balanced salt solution (HBSS, pH 7.4) for 15 minutes at 37°C. Cells were then washed thrice with PBS, fixed using 4% para-formaldehyde (PFA), counterstained with Vecta-shield DAPI and mounted onto glass slides. Images were acquired on the LSM 780 confocal microscope.

#### Flow cytometry

Cells were stained with MitoTracker Red CMXRos (100nM) diluted in 2ml of Hank’s balanced salt solution (HBSS, pH 7.4) for 15 minutes at 37°C. Cells were washed twice with pre-warm HBSS, collected in FACs tubes using cell scraper. Samples were acquired on Attune NxT flow cytometer (ThermoFisher Scientific, USA).

### Assessment of mitochondrial membrane protein TOM20

NIH3T3 cells were treated with cfChPs (10 ng) for 4 h and immunostaining for TOM20 was performed by fixing the cells with 4% PFA for 15 minutes at 37°C followed by permeabilization with 0.2% Triton X-100 for 30 min and blocking with 3% BSA for 1 h. Cells were immune-stained using anti-rabbit TOM20 primary antibody and appropriate secondary anti-rabbit FITC antibody followed by mounting with Vecta-shield DAPI. Images were acquired on the Leica confocal SP8 imaging system.

### Assessing mitochondrial membrane potential using JC-1 dye

#### Fluorescence microscopy

After treatment with cfChPs (10 ng) for 4 h, NIH3T3 cells were stained with JC-1 dye in HBSS and incubated at 37°C for 15 min. The cells were washed twice with pre-warmed HBSS. Cells were counterstained with Hoechst, mounted on glass slides and cell-associated fluorescence was detected using Spectral Bio-Imaging System (Applied Spectral Imaging).

#### Flow cytometry

Cells were stained with JC-1 dye in HBSS and incubated at 37°C for 15 min. Cells were washed twice with pre-warm HBSS, collected in FACs tubes using cell scraper. Samples were acquired on Attune NxT flow cytometer (ThermoFisher Scientific, USA).

### Assessment of mitochondrial ROS production using MitoSOX Red

#### Fluorescence microscopy

NIH3T3 cells were treated with cfChPs (10 ng) for 4 h and stained with 0.5 µM MitoSOX Red. Cells were in 2 ml of HBSS for 15 minutes at 37°C, washed with warm buffer. Cells were counter stained with Hoechst, mounted on glass slides and cell-associated fluorescence was measured using Spectral Bio-Imaging System (Applied Spectral Imaging).

#### Flow cytometry

Cells were stained with 0.5 µM MitoSOX Red diluted in 2 ml of HBSS for 15 minutes at 37°C, washed with pre-warm HBSS and then were collected in FACs tubes using cell scraper. Samples were acquired on Attune NxT flow cytometer (ThermoFisher Scientific, USA).

### Assessment of superoxide dismutase (SOD-1) expression

#### Fluorescence microscopy

cfChPs treated NIH3T3 cells were fixed with 4% paraformaldehyde (PFA) for 15 minutes at 37°C and were then immune-stained using primary antibody against SOD-1 and appropriate secondary antibody according to the protocol described above. Cells were mounted with Vecta-shield DAPI, and images were acquired using Spectral Bio-Imaging System (Applied Spectral Imaging).

#### Flow cytometry

Cells were fixed with 4% paraformaldehyde (PFA) for 15 minutes at 37°C and were then stained using primary antibody against SOD-1 and appropriate secondary antibody. Cells were acquired on Attune NxT flow cytometer (ThermoFisher Scientific, USA).

### Procedure for collecting conditioned media from hypoxia induced dying NIH3T3 cells

A dual chamber system was used to generate culture medium containing cfChPs released from dying cells. NIH3T3 cells were seeded at a density of 2 x 10^5^ on ThinCert^®^ Cell Culture Inserts (pore size 400 nm) containing 1.5 ml of DMEM and were placed in a six-well culture plate. Cells were incubated at 37 °C for 24 h. The six-well plate with Thincert^®^ Inserts was then transferred to a hypoxia chamber with 1% O_2_ to induce hypoxic cell death. After 48h, the plate was removed from the hypoxia chamber and the 700 μl of DMEM media was added to the lower chamber of the six-well palte below the ThinCert^®^ Inserts and kept for 48 h at 37°C in normoxic conditions to allow cfChPs <400 nm in size to seep into the medium in the lower chamber. The medium in the lower chamber was collected and divided into aliquots and stored at -80°C until further use.

### Assessment of mitochondrial ROS production using conditioned medium

Aliquots were thawed for use in the dose / volume response experiments which involved addition of increasing volumes of the culture medium to NIH3T3 cells, and the level of ROS was measured by MitoSOX Red staining. A dose / volume of 50 µl was found to be the most effective in generating ROS (Supplementary Fig. 1). Once the optimal dose / volume was ascertained, aliquots were thawed for experiments involving the use of three different cfChPs deactivators viz. pullulan-histone antibody nanoconjugates (CNPs), DNase I and a novel pro-oxidant combination of small quantities of resveratrol and copper (R-Cu) (please see below).

### Pullulan-histone antibody nanoconjugates (CNPs)

Pullulan-histone antibody nanoconjugates (complexed nanoparticles, CNPs) were synthesized according to the protocol described by us earlier^39^.

### DNase I

DNase I from bovine pancreas was procured from Sigma-Aldrich (Catalogue No. DN25-1G).

### Resveratrol-copper (R-Cu)

Resveratrol (R) is a plant polyphenol with antioxidant properties^40^. However, R acts as a pro-oxidant in the presence of copper (Cu) by reducing Cu (II) to Cu (I) thereby generating a free radical^41, 42^. R-Cu can deactivate cfChPs in *in vitro*^19, 34^. The involved concentrations of R and Cu used in this study were 1mM and 0.0001mM respectively.

### Experiments using conditioned medium with or without cfChPs deactivators

Conditioned media were pre-treated with the above three cfChPs deactivators for 1h at 37°C prior to addition to NIH3T3 cells. ROS production in NIH3T3 cells were measured at 4h using MitoSox Red (protocol mentioned above).

### Statistical analysis

All data are presented as Mean_±_Standard Error of Mean (SEM). Statistical analysis was performed using GraphPad Prism 8 (GraphPad Software, Inc., USA, Version 8.0). Data were compared using Student’s_t*-*_test (two tailed, unpaired) and one-way ANNOVA. *p*_<_0.05 was taken as the level of significance.

### Data Availability

All data are contained within the manuscript. Any additional data will be provided on reasonable request.

## Legends to supplementary figures

**S1 Fig.** Representative fluorescence microscopy images showing bystander uptake of dually labelled cfChPs by NIH3T3 cells at 4h. Donor NIH3T3 cells were dually labelled in their DNA with BrdU (red) and histones by CellLight® Histone 2B-GFP (green). Cells were kept in hypoxic chamber for 48h and the conditioned medium (50µl) containing dually labelled cfChPs were added to healthy recipient NIH3T3 cells.

**S2 Fig.** Dose / volume response analysis of activation of ROS production as detected by MitoSOX at 4h. NIH3T3 cells treated with increasing volume of conditioned medium containing cfChPs released from hypoxia induced dying NIH3T3 cells. Maximum activation of MitoSOX is seen with 50µl of culture medium.

**S3 Fig.** Graphical illustration showing cfChPs that circulate in blood (left hand image) and those that are released locally from dying cells (right hand image) can readily enter into healthy cells to damage their mitochondria leading to ROS production.

## Acknowledgement

The authors thank Mr. Ashish Pawar for his help in preparing manuscript.

## Author contributions

G.V.R., B.K.T., K.A., S.S., N.K.K., R.L., K.P. conducted the experiment; G.V.R., B.K.T., K.A., S.S. conducted data analysis; G.V.R. supervised the experiments; G.V.R., B.K.T., K.A., S.S., I.M. wrote the paper; I.M. conceived the idea, procured funding, supervised the experiments, wrote the paper and approved the final manuscript.

