## Supplementary figures and images for "Cell-free chromatin particles released from dying cells inflict mitochondrial damage and ROS production in living cells"

### Supplementary Figure 1

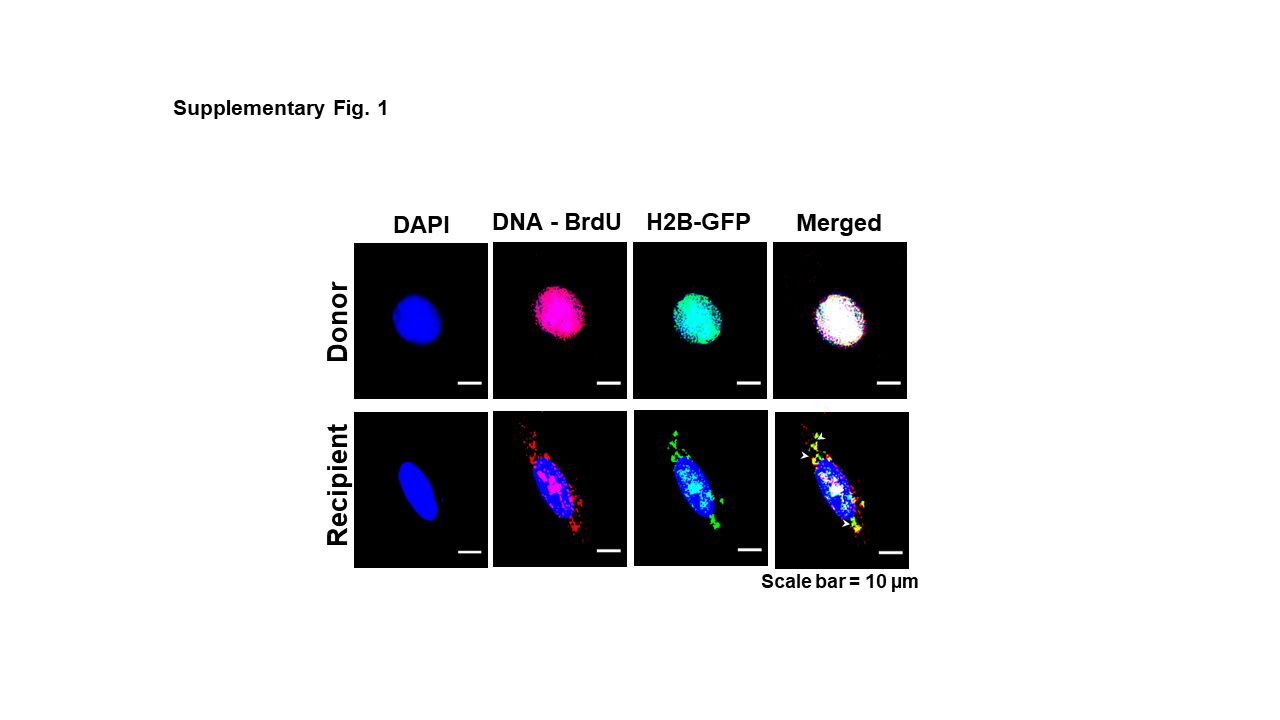

### Supplementary Figure 2

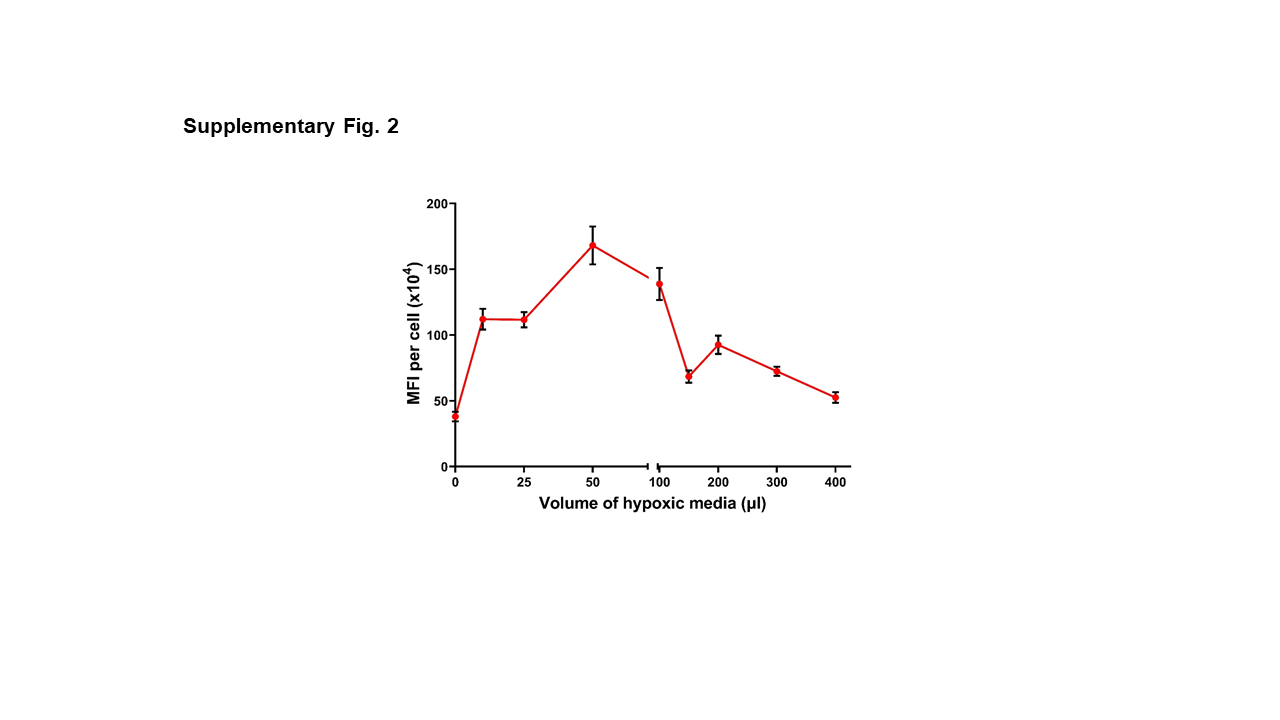

### Supplementary Figure 3

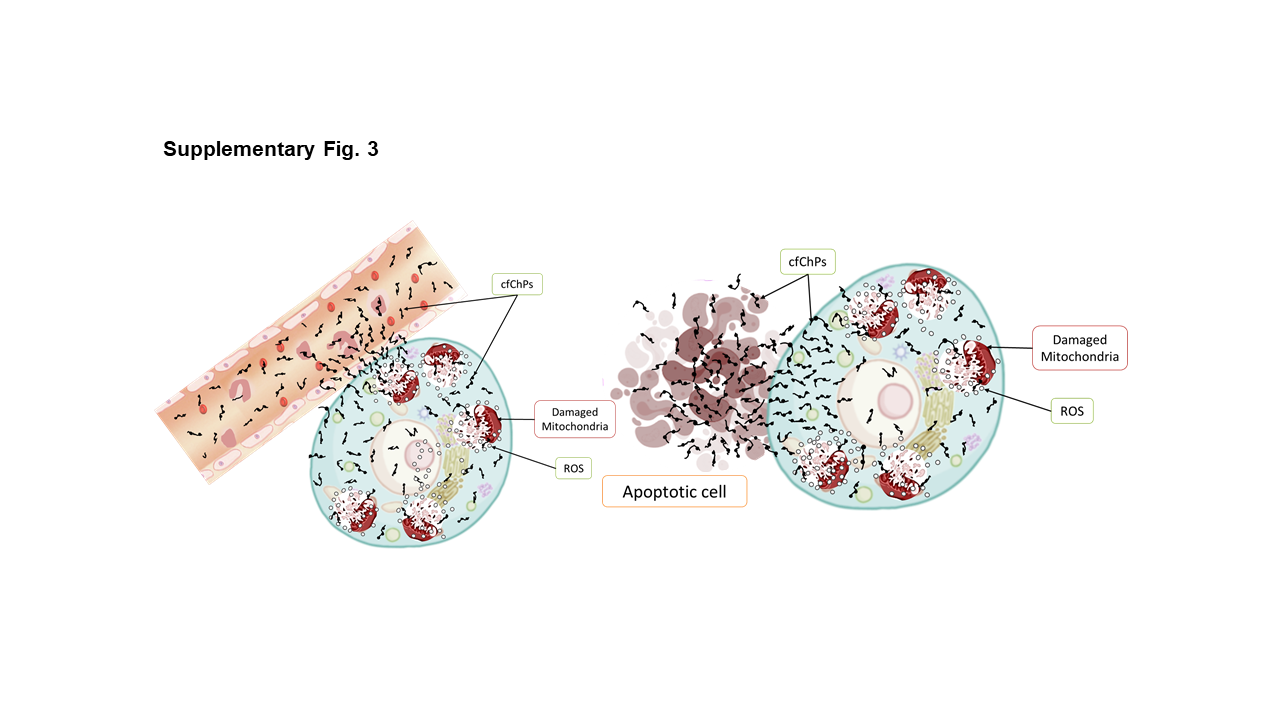
