## Supplementary Table 1 for "Cell-free chromatin particles released from dying cells inflict mitochondrial damage and ROS production in living cells"

**Sources of reagents and antibodies used in this study.**

List of reagents used.

| **Sl. No.** | **Reagent** | **Catalogue Number** | **Company** |
| --- | --- | --- | --- |
| 1 | Platinum Bright 550 Red | GLK-004 | Kreatech Diagnostics, Denmark |
| 2 | ATTO 488 | AD 488 | ATTO-TEC, GmbH, Germany |
| 3 | MitoSOX Red | M36008 | Thermo Fisher Scientific, USA |
| 4 | MitoTracker Green FM | M7514 | Thermo Fisher Scientific, USA |
| 5 | MitoTracker Red CMX Ros | M7512 | Thermo Fisher Scientific, USA |
| 6 | Hoechst 33342 | H21492 | Thermo Fisher Scientific, USA |
| 7 | Vecta-Shield DAPI | 101098-042 | Vector Laboratories, USA |
| 8 | MitoScreen kit (JC-1) | 551302 | BD Biosciences, USA |
| 9 | Mitochondria Isolation Kit | 89874 | Thermo Fisher Scientific, USA |

List of antibodies used

| **Sl. No.** | **Antibody** | **Catalogue Number** | **Company** |
| --- | --- | --- | --- |
| 1 | Rabbit monoclonal anti- TOMM20 antibody | ab186735 | Abcam, United Kingdom |
| 2 | Rabbit monoclonal anti-Superoxide Dismutase-1 antibody | ab51254 | Abcam, United Kingdom |
| 3 | Mouse monoclonal anti- γ-H2AX | 05-636 | Merck, GmbH, Germany |
| 4 | Mouse monoclonal anti-phospho-ATM (Ser 1981) antibody | 05-740 | Merck, GmbH, Germany |
| 5 | Goat Anti-Rabbit IgG H&L (FITC) | ab6717 | Abcam, United Kingdom |
| 7 | Rabbit Anti-mouse (FITC) | AP160F | Merck, GmbH, Germany |
